# ERβ mediates sex-specific protection in the *App-NL-G-F* mouse model of Alzheimer’s disease

**DOI:** 10.1101/2024.07.22.604543

**Authors:** Aphrodite Demetriou, Birgitta Lindqvist, Heba G. Ali, Mohamed M. Shamekh, Silvia Maioli, Jose Inzunza, Mukesh Varshney, Per Nilsson, Ivan Nalvarte

## Abstract

Menopausal loss of neuroprotective estrogen is thought to contribute to the sex differences in Alzheimer’s disease (AD). Activation of estrogen receptor beta (ERβ) can be clinically relevant since it avoids the negative systemic effects of ERα activation. However, very few studies have explored ERβ-mediated neuroprotection in AD, and no information on its contribution to the sex differences in AD exists. In the present study we specifically explored the role of ERβ in mediating sex-specific protection against AD pathology in the clinically relevant *App^NL-G-F^* knock-in mouse model of amyloidosis, and if surgical menopause (ovariectomy) modulates pathology in this model. We treated male and female *App^NL-G-F^* mice with the selective ERβ agonist LY500307 and subset of the females was ovariectomized prior to treatment. Memory performance was assessed and a battery of biochemical assays were used to evaluate amyloid pathology and neuroinflammation. Primary microglial cultures from male and female wild-type and ERβ-knockout mice were used to assess ERβ’s effect on microglial activation and phagocytosis. We find that ERβ activation protects against amyloid pathology and cognitive decline in male and female *App^NL-G-F^* mice. Ovariectomy increased soluble amyloid beta (Aβ) in cortex and insoluble Aβ in hippocampus, but had otherwise limited effects on pathology. We further identify that ERβ does not alter APP processing, but rather exerts its protection through amyloid scavenging that at least in part is mediated via microglia in a sex-specific manner. Combined, we provide new understanding to the sex differences in AD by demonstrating that ERβ protects against AD pathology differently in males and females, warranting reassessment of ERβ in combating AD.

## 1. INTRODUCTION

Over recent years, increasing number of studies have suggested that the female sex hormone estrogen (E2) elicit neuroprotective functions, which are lost upon menopause, and that this loss may at least partly account for the increased female prevalence to Alzheimer’s disease (AD) (1-3). Indeed, bilateral oophorectomy has been identified as a possible risk factor for dementia (4-6). However, since all aged women enter menopause, but not all get dementia, other risk factors must exist that interact with lower circulating E2 levels.

Three types of estrogen receptors are found in the brain, estrogen receptor alpha (ERα), beta (ERβ), and the G-protein coupled estrogen receptor (GPER1). While ERα is highly expressed in hypothalamus to regulate functions related to reproduction, the roles of GPER1 and ERβ are less clear, and all three receptors are expressed in regions important for cognitive behavior such as the cortex and hippocampus (1). Previous studies have proposed ERβ to be of particular interest as a possible therapeutic target in mediating neuroprotection since its activation is, unlike that of ERα, not associated with adverse health effects (7). In the context of AD, ERβ has been suggested to play multifaceted role in neuroprotection and neuronal survival (8-12). However, variation between studies have led to inconclusiveness and there is a gap in knowledge to the exact contribution of ERβ to the sex-differences in AD. Many plausible explanations could be the causes, including the usage of different AD mouse models with no direct comparison between the sexes. In addition, different types of ERβ ligands with varying selectivity have been used with different results, adding to inconclusiveness.

In this study, we focus selectively on the role of ERβ in mitigating amyloid pathology in the *App^NL-G-F^* knock-in mouse model that exhibits robust Aβ pathology (but without APP overexpression), neuroinflammation, synaptic alterations, and behavior impairment (13). We evaluate the effect of the highly potent selective ERβ agonist LY500307 on AD pathology in male and female *App^NL-G-F^* mice. LY500307 has passed first lines of toxicity and safety tests and is currently in phase 2 clinical trials for alleviation of perimenopausal depression (Clinical trials identifier: NCT03689543), rendering it a human-relevant drug to test in our chosen AD model. Our data show that LY500307 protects against Aβ plaque buildup in cortex and hippocampus, as well as against cognitive deterioration in both male and female *App^NL-G-F^* mice. Although ERβ activation does not affect APP processing, it reduces number of activated microglia in both males and females, but with stronger effects in males. We also show that loss of ERβ in microglia modulates microglial activation and amyloid scavenging in a sex-specific manner. Finally, we show that removal of systemic E2 levels by ovariectomy (surgical menopause) has limited effects on amyloidosis in *App^NL-G-F^* females. Our data contribute to the increased understanding to the sex differences in AD and warrants further exploration of ERβ as a potential therapeutic target in AD.

## 2. MATERIAL AND METHODS

### 2.1 Animals and treatments

Male and female *APP^NL-G-F^* knock-in mice (carrying the Swedish [NL], Arctic [G] and Iberian [F] mutations in the humanized Aβ peptide (13)) were obtained from local breeding using the C57/BL6J strain background. At 2.5 months of age, female mice were selected randomly for bilateral ovariectomy or for sham surgery. Similarly, at 3 months of age male and female mice were randomly selected for LY500307 (0.35mg/kg/day, Santa Cruz Biotechnology, Dallas, TX, USA), dissolved in vehicle solution (40% Captisol, [Cydex pharmaceuticals, Lawrence, KS, USA], 1% ethanol, and 59% 0.1M PBS), or vehicle treatment (vehicle solution) through oral gavage administration. The treatment regimen was daily delivery over 7 days, followed by 7 days of rest. The resting period was included to avoid hormone-induced downregulation of ERβ gene (14). This was repeated twice after which animals were subjected to behavior studies (2 days after last treatment and 2 days of rest between tests) and sacrificed at 5 months of age. For brain dissection, animals were deeply anesthetized with isoflurane followed by intracardial ice-cold 0.1M PBS perfusion. Half brain was fixed in cold 4% paraformaldehyde and hippocampus and cerebral cortex of the other half was snap-frozen for biochemical assays. All procedures were performed in accordance with approved ethical permits (ethical approval ID 407 and ID 2199-2021, Linköping’s animal ethical board).

### 2.2. Behavioral tests

#### 2.2.1. Contextual cued fear conditioning

A conditioning semi-transparent plexiglass chamber of 17x17x25 cm (l x w x h) with a stainless-steel grid floor (grid spaced 0.5cm apart, Ugo Basile, Gemonio, Italy) surrounded by sound-attenuating grey chest was used for training and conditioning tests under a constant light (50 Lux) and background white noise (77db). The chest was fitted with a light sensitive camera over the chamber. The chamber was cleaned with 70% ethanol before each individual mouse tests. The contextual fear conditioning test was performed over a span of 3 days, as previously described (15). Briefly, on the conditioning day, mice were individually and randomly placed in the chamber and allowed to explore for 2 min before the onset of the conditional stimuli in the form of two sound exposures (65db, 2000 Hz) 1 min apart lasting for 30 s each. During the last 2 s of each conditioning stimulus the mice received a mild electric foot shock (0.5 mA). The conditioning ended 1 min after the last shock. The next day, the mice were subjected to the contextual test where they were placed back in the chamber for 3 min but were not subjected to any sound stimuli or foot shock. On the third day, the mice were subjected to the cued test in which they were placed back in the chamber that had been fitted with different environment (checkered wall patterns and white bottom). The mice were free to explore the chamber for 2 min (baseline) before the onset of the sound stimuli (cue tone, 65db, 2000 Hz) for the rest 2 min without any foot shock. Mouse movement was traced by a computer-based video tracking system (ANY-Maze 6.3 software, Stoelting, Dublin, Ireland). The freezing response was defined as the percentage of time a mouse remained motionless (divided into 30 sec intervals).

#### 2.2.2. Y-maze

Spatial working memory and reference memory were analyzed using the standard Y-maze test. The Y-maze was used for hippocampal dependent spontaneous alteration measurements, and consisted of 3 arms (35 x 7 x 15 cm, made of non-reflective gray plastic, Noldus Wageningen, Netherlands) at 120° angle to each other. A random mouse from each test group was placed in the center of the maze, and the 5 min trial started when the experimenter was out of the room of the maze to allow uninterrupted movement of the animal. Both manual and automated recording (using EthoVision XT, Noldus, Wageningen, Netherlands) of numbers of entries into each arm was used to calculate the percent spontaneous alterations. Alternations were considered to be completed when a mouse performed successive entries into three different arms. Percentage alternations was calculated as the ratio of actual alternations and maximum possible alternations. The Y-maze was cleaned with 70% ethanol before each individual mouse tests.

### 2.3. Immunohistochemical and histochemical analyses

4µm thick paraffin-embedded sagittal mouse brain sections were fixed on glass slides, hydrated, followed by heat-induced antigen retrieval in a pressure steamer at 121°C for 20 min, followed by 15 min permeabilization with 0.5% Triton-X 100 (Millipore, Burlington, MA, USA) and blocking using 10% Horse Serum (ThermoFisher Scientific, Waltham, MA, USA), 0.1% Tween-20 (Millipore) in 0.1M PBS for 1h at 37°C. Following blocking the slides were immunostained over-night at 4°C with antibodies specific to Aβ (1:2000 dilution, 82E1, IBL-Tecan, Männedorf, Switzerland), Iba1 (1:300, ab178846; and 1:300 ab225260, both from Abcam, Cambridge, UK), GFAP (1:300, GA5 Alexa Fluor_488_-labeled, Millipore), CD68 (1:300, Ab283654, Abcam), and/or ERβ (1:5000, PP-PPZ0506-00, R&D Systems, Minneapolis, MN, USA) (Supplemental Table 1). For antibodies raised in mice, we used 1x mouse-on-mouse IgG blocking solution (ThermoFisher Scientific) prior to antibody incubation. Secondary antibodies were Alexa Fluor_488_, Alexa Fluor_568_ (both from ThemoFisher Scientific). To reduce autofluorescence, the sections were incubated in 1 mM CuSO_4_ diluted in 50 mM ammonium acetate for 15 min. Nuclear staining was with 300 nM DAPI (ThermoFischer Scientific) for 10 min, prior to mounting. To visualize amyloid plaques, we used 1x AmyloGlo stain (Biosensis, Thebarton, Australia) supplemented to the secondary antibody solution. ABC-HRP kit and Impact-DAB (both from Vector Laboratories, Newark, CA, USA) were used for immunohistochemical staining according to manufacturer’s recommendations. Immunofluorescence images were captured using an AxioPlan-2 fluorescent microscope (Carl Zeiss, Oberkochen, Germany) and the Zeiss AxioVision 4.0 software (Carl Zeiss). Image analysis was performed on at least 3 sections per mouse using the ImageJ software (NIH, Bethesda, MD, USA) and setting image threshold and counting was as described previously (16). For each acquired image, the image lookup table (LUT) was kept linear and covered the whole image data. Association of microglia to plaques were quantified by counting number of microglia within 20 µm radius of plaque edge.

### 2.4. Aβ and cytokine profile immunoassays

Frozen cortical and hippocampal tissues were thawed and homogenized in ice-cold TBS buffer (50 mM Tris-HCl, pH 7.6, 150 mM NaCl, and protease inhibitor cocktail (Roche, Basel, Switzerland)). The homogenates were centrifuged at 24 000 x g for 45 min at +4°C, yielding a soluble fraction (supernatant) and an insoluble fraction (pellet). The pellets were solubilized by resuspension in 6M Guanidine-HCl and sonication using a water-bath sonicator (Bioruptor, 5 min max output (Diagenode, Denville, NJ, USA)). Soluble pellets were centrifuged at 24 000 x g for 45 min at +4°C and the supernatant (defined as insoluble fraction) was diluted in TBS to yield 0.5 M Guanidine-HCl. Similarly, Guanidine-HCl was added to the soluble fractions to yield a concentration of 0.5 M Guanidine-HCl. Total protein concentration was determined using the BCA Protein Assay (ThermoFisher Scientific) or the Coomassie Protein Assay reagent (Sigma-Aldrich, St Louis, MO, USA). Quantification of Aβ_1-40_ and Aβ_1-42_ in soluble and insoluble fractions was performed using the EZHS-SET ELISA kit, following manufacturer’s instructions (Millipore) and read on a Tecan plate spectrophotometer. Proinflammatory cytokine profiling on soluble fractions were performed using the V-PLEX proinflammatory panel 1 (mouse) kit (Mesoscale Discovery) on soluble brain fractions according to manufacturer’s instructions. The kit allows multiplex quantification of IFN-γ, IL-1β, IL-2, IL-4, IL-5, IL-6, IL-10, CXCL1 (KC/GRO, keratinocyte-derived chemokine/growth-related oncogene), IL12p70, and TNF-α. Samples were read on the MESO QuickPlex SQ120 reader and data were analyzed using the Discovery Workbench 4.0 software (both from Mesoscale Discovery). The concentration of each cytokine in the tissue lysates was normalized with the total protein concentration of the respective sample.

### 2.5. Western blot

Cortical and hippocampal tissue were homogenized in ice-cold 4x PIPES buffer pH 6.8 (40 mM Piperazine-1,4-bis(2-ethanesulfonic acid), 1.2M Sucrose, 0.4M NaCl, 27 mM MgCl_2_, and 1x protease inhibitor cocktail, all from Sigma). Cell debris were pelleted and supernatant were centrifuged at 24 000 x g for 45 min at +4°C. The pellet was resuspended in a low volume of 4x PIPIES buffer and protein concentration was measured and adjusted to 2.5 mg/ml. 100µg protein were incubated at 37°C for 30 min followed by chloroform-methanol protein precipitation. In brief, 600µl of cholorform:methanol in ratio 2:1 was added to the protein mixture and incubated for 30 min at room temperature (RT) under agitation followed by centrifugation and phase separation at 24 000 x g for 15 min at RT. The intermediate was isolated, resuspended in 600µl cholorform:methanol 1:2 and incubated for 60 min at RT under agitation. The protein was precipitated by centrifugation at 24 000 x g for 15 min at RT, supernatant was removed, and pellet was let to dry. The protein pellet was resuspended in SDS-loading buffer to yield 3mg/ml. In brief, 10-30µg of protein were loaded on 4-20% gradient SDS-PAGE gels and proteins were transferred to a PVDF membrane. After blocking the membrane was subjected to antibody against Aβ1-16 (6E10, BioLegend), APP N-terminus (22C11, Millipore), APP C-terminus (A8717, Sigma-Aldrich) and antibody against β-Actin (AC-15, Millipore). Detection was performed using ECL substrate (ThermoFisher) and exposure to light-sensitive films or CCD camera. Quantification of bands was performed using ImageJ software (NIH). All blots were processed in parallel.

### 2.6. Real-time quantitative PCR analysis

Total RNA from cells or tissue was extracted using the RNeasy plus mini kit, RNeasy plus micro kit or Allprep DNA/RNA kit (Qiagen) according to manufacturer’s instructions, and RNA concentrations and quality were determined with NanoDrop (ThermoFisher Scientific). Complementary DNA was synthesized using SuperScript IV VILO Master Mix cDNA synthesis kit (ThermoFisher Scientific). The qPCR reaction contained 5 or 10 ng of cDNA, exon-exon spanning primers (500 nM), and KAPA SYBR Fast qPCR master mix (Sigma-Aldrich) or using TaqMan assays (Supplemental Table 2) and TaqMan Fast Advanced Master Mix (Applied Biosystems) and was performed on an ABI 7500 fast thermal cycler (Applied Biosystems) according to manufacturer’s instructions. Expression relative to housekeeping gene was calculated using the ΔCt method.

### 2.7. Primary microglial isolation from adult mouse brain and culture

Primary mixed neuroglia cultures were prepared from adult male and female WT and all-exon *Esr2* deleted (*Esr2*-KO) C57BL/6NTac mice (17) as described previously (18). Briefly, adult mice (6 months old) were terminally transcardially perfused with cold PBS under isoflurane inhalation anesthesia. Meninges were removed and the brains were collected in ice-cold Hibernate-A medium. Brain tissues were gently mechanically triturated, and enzymatic digestion was followed in 0.25% Trypsin (without phenol red) with 200 U/mL DNase I (Gibco/ThermoFisher Scientific) at RT for 30 minutes on orbital shaker. Sequentially, an equal volume of Defined Trypsin Inhibitor (Gibco/ThermoFisher Scientific) was added, and the homogenate was passed through 100 μm and 40 μm pre-wetted cell strainer (VWR) to form a single-cell suspension. The single-cell suspension was cultured in a poly-D-lysine coated T75 flask in DMEM-F12 medium (without phenol red) containing 10% heat-inactivated fetal bovine serum (FBS), microglia growth supplement (MGS, ScienCell, Carlsbad CA, USA) and antibiotic/antimycotic (ThermoFisher Scientific) for 15 to 28 days in 5% CO_2_ at 37°C. Culture medium was changed every three days. Microglia were separated from mixed glial cell cultures via magnetic separation using MiniMACS® CD146 magnetic bead negative selection (to remove endothelial cells) and CD11b magnetic bead positive selection according to the manufacturer’s protocols (Miltenyi Biotech, Bergisch Gladbach, Germany). Briefly, cells were dissociated with Accumax (Gibco/ThermoFisher Scientific), washed, and incubated with FcR blocking reagent (Miltenyi Biotech) followed by CD146 microbeads in FACS buffer (1% FBS in DPBS). The negative fraction was further incubated with CD11b microbeads and CD11b positive cells were quantified using an automated cell counter (Countess II FL, ThermoFischer Scientific). Primary microglia were cultured as described above without MGS, for further analyses under resting or stimulated conditions.

### 2.8. Microglia stimulation

The cells were treated with LY500307 (10 nM, Santa Cruz) or vehicle (ethanol) for 24 h prior to stimulation and then additionally treated for 24 h with lipopolysaccharide (100 ng/mL, LPS, Sigma-Aldrich) and IFN-γ (20 ng/mL) (M1 stimulation) or IL-4 (20 ng/mL) and IL-13 (15 ng/mL, all from Peprotech) (M2 stimulation).

### 2.9. Phagocytosis assay

Fluorescent Amyloid beta (Aβ_1-42_) peptide (HyliteFluor 488, Anaspec, Fremont, CA, USA), 205 nM in DMSO was incubated with M1, M2 or control microglia for 24 h at 37°C in 5% CO_2_. The cells were washed thoroughly and the uptake of Aβ was analyzed using SpectraMax i3x Imaging Cytometer (Molecular Devices). Unstimulated cells incubated with Aβ were used as control, cells without added Aβ served as negative control (subtracted from all values) while cells incubated with the phagocytosis inhibitor Cytochalasin D (20 nM in DMSO, ThermoFisher Scientific) 2 hours prior Aβ incubation inhibited phagocytosis and was used for normalization of the data.

### 2.10. Migration assay

The chemotactic migration of microglia was assessed in a 24-well Millicell cell culture insert plate (8 μm pore diameter, Millipore). The upper chambers of the wells were loaded with cells and incubated for 1 h at 37 °C in 5% CO_2_. The cells were treated with stimulants as mentioned above. The following day, chemoattractant CX3CL1 (20 ng/ml Fractalkine, R&D systems, Minneapolis, MN, USA) was added to the lower chamber while negative control contained 0 ng/ml chemoattractant and positive control contained 20 ng/ml to both upper and lower chamber. Once loaded, cells were incubated for 5 hours at 37°C and 5% CO_2_. Following incubation, the inserts were thoroughly rinsed in PBS to remove all cell debris and then fixed in 4% paraformaldehyde for 15 min, followed by permeabilization with 0.2% Triton X-100 in PBS and stained with DAPI for 10 min. The non-migrated cells were removed from the top side of the filter whilst cells that migrated toward the bottom of the filter were quantified. To quantify the migrated cells, several non-overlapping images were captured before and after cleaning the top side of the filter for each well/condition and processed in Fiji (ImageJ, NIH, Bethesda, MD, USA) for manual nuclei counting.

### 2.11. Statistical analysis

Results are expressed as means ±SD. The statistical analyses were performed using GraphPad Prism 9.02 software (GraphPad Software, San Diego, USA). Data were tested for equal variance by *F*-tests. Unpaired two-tailed Student’s t tests were used to compare between two groups. Unless stated otherwise, multiple group analyses were performed by two-way or three-way analysis of variance (ANOVA), followed by uncorrected Fisher’s LSD test or corrected post-hoc tests for multiple comparisons as indicated in figure legends. Significance level was set at <0.05 (\**P* < 0.05, \*\**P* < 0.01, \*\*\**P* < 0.001, \*\*\*\**P* < 0.0001). All analyses are based on at least 3 biological replicates.

## 3. RESULTS

### 3.1. ERβ expression in the mouse cortex and hippocampus

Since estrogen (E2) has been ascribed neuroprotective properties (1-3), we sought to explore if selective activation of the estrogen receptor beta (ERβ, *Esr2* gene product), a more clinically relevant target than ERα, can be protective against amyloid-related pathology in the *App^NL-G-F^* mouse model of AD. Since expression of ERβ in the brain has been questioned due to poor antibody specificities to ERβ, we first performed immunohistochemical analysis using a validated ERβ antibody in wild-type (WT) and ERβ knockout (*Esr2*-KO) mouse brains. This revealed scattered expression with both cytoplasmic nuclear localization in several brain regions affected in AD, including the frontotemporal, primary motor, somatosensory and visual cortices, as well as in the granule layers of CA2 and dentate gyrus (DG) of the hippocampus (Supplemental Fig. 1A, B). Highest number of ERβ positive cells were seen in frontal and primary motor cortex as well as in the hippocampus (Supplemental Fig. 1B). ERα (*Esr1*) and Erβ (*Esr2*) had similar expression between male and female mice in cortex and hippocampus although ERα expression was about 5-10 fold higher than ERβ in both brain regions, which did not change upon ERβ loss (Supplemental Fig. 1C-F).

### 3.2. ERβ activation protects against cognitive deficits and Aβ_42_ deposition in *App^NL-G-F^* mice

To study effect of ERβ activation on AD pathology we treated *App^NL-G-F^* male and female mice with the selective ERβ agonist LY500307 (with 12-fold higher selectivity for ERβ over ERα and 32-fold more functional potency (19, 20)), 0.35 mg/kg daily through oral gavage every other week over 5 weeks, starting at 3 months of age. A subset of female mice was ovariectomized (OVX) 2 weeks prior to treatment to study the influence of loss of circulating E2 (Fig. 1A). At the end of the treatment, the mice were subjected to memory tests, where the mice treated with LY500307 (LY) performed better than vehicle treated mice in the Y-maze spatial memory test, though we did not see any effect of OVX (Fig. 1B). Similarly, mice given LY performed better than vehicle treated mice in the cued fear conditioning (FC) associative memory test where LY treated mice had longer episodes of freezing both during the contextual (Fig. 1C) and cued tests (Fig. 1D, E). Interestingly, LY treated females also displayed increased freezing at baseline, even in the new context, suggestive of an overall better memory performance after LY treatment (Fig 1E). OVX mice did not have a negative effect on memory performance, oppositely, it slightly improved memory in the contextual FC test although less significant compared to LY treatment (Fig. 1C). Overall, these results suggest that ERβ activation may indeed act neuroprotective in the *App^NL-G-F^* model.

**FIGURE 1.**
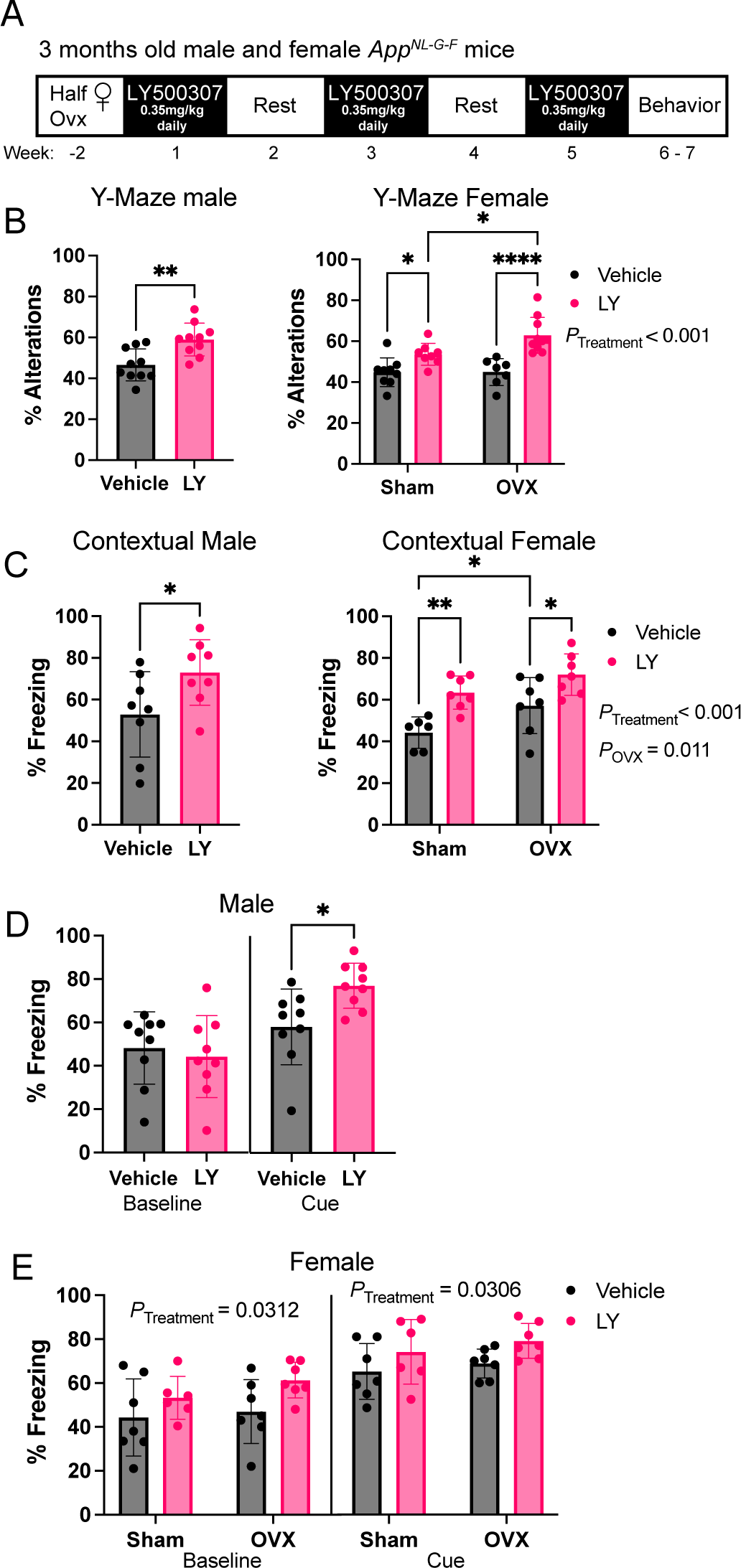
ERβ activation improves cognitive behavior in *App^NL-G-F^* male and female mice. (A) Treatment regime of *App^NL-G-F^* mice. (B) Percent Y-maze alterations of male (left) and female (right) *App^NL-G-F^* mice treated with vehicle or ERβ agonist LY500307 (LY) (n = 7-10). (C) Percent context-associated freezing time of male (left) and female (right) *App^NL-G-F^* mice (n = 6-9) in the contextual fear conditioning test. Cued-associated freezing time in the contextual fear conditioning test of (D) male and (E) female *App^NL-G-F^* mice before cue (baseline) and upon cue (tone) in a different cage context (n = 6-9). Female mice were either ovariectomized (OVX) or sham operated (Sham). * *P* < 0.05, ** *P* < 0.01, *** *P* < 0.001. Unpaired t-test was used for males and 2-way ANOVA for females followed by uncorrected Fisher’s LSD test for multiple comparisons. Overall significant main effects of treatment or OVX are indicated.

Next, we analyzed the effect of LY on amyloid pathology. LY treated mice had generally lower number and smaller size of amyloid plaques in different cortical regions and in hippocampus in both male (Fig. 2A-C) and female (Fig. 2D-F) *App^NL-G-F^* mice. When taking number of ERβ positive cells into account in those regions, we could observe that the largest effect size of LY treatment in lowering Aβ plaques overlapped with highest levels of ERβ positive cells in those regions (Fig. 2G, Supplemental Fig. 1B). OVX did not have any effect on number of plaques, but slightly (but significantly) increased existing plaque area in visual and somatosensory (Vis/Ss) cortex (Fig. 2E-F). In line with these results, the levels of soluble and insoluble neurotoxic amyloid beta (Aβ_42_) were overall lower in cortex and hippocampus in both male (Fig. 2H, I) and female (Fig. 2J, K) *App^NL-G-F^* mice after LY treatment, although it did not reach statistical significance for hippocampal soluble and cortical insoluble Aβ_42_ in male mice and no statistical significance for cortical insoluble Aβ_42_ levels in females. Interestingly, OVX increased Aβ_42_ levels in the cortex, while having no effects on Aβ_42_ levels in other brain areas. Aβ_40_ levels were similar to Aβ_42_ levels (Supplemental Fig. 2A-D) and Aβ_42_/Aβ_40_ ratio did not differ with LY treatment, although there was an overall increase in soluble Aβ_42_/Aβ_40_ ratio in female cortex upon OVX (Supplemental Fig. 2F). Furthermore, female mice had in general higher levels of soluble Aβ_42_ levels compared to male mice (Supplemental Fig. 2G, H). These data suggest that ERβ activation reduces Aβ levels and plaque load in both male and female *App^NL-G-F^* mice, but sex differences exist, and that OVX can worsen amyloid pathology although differently in in different brain regions.

**FIGURE 2.**
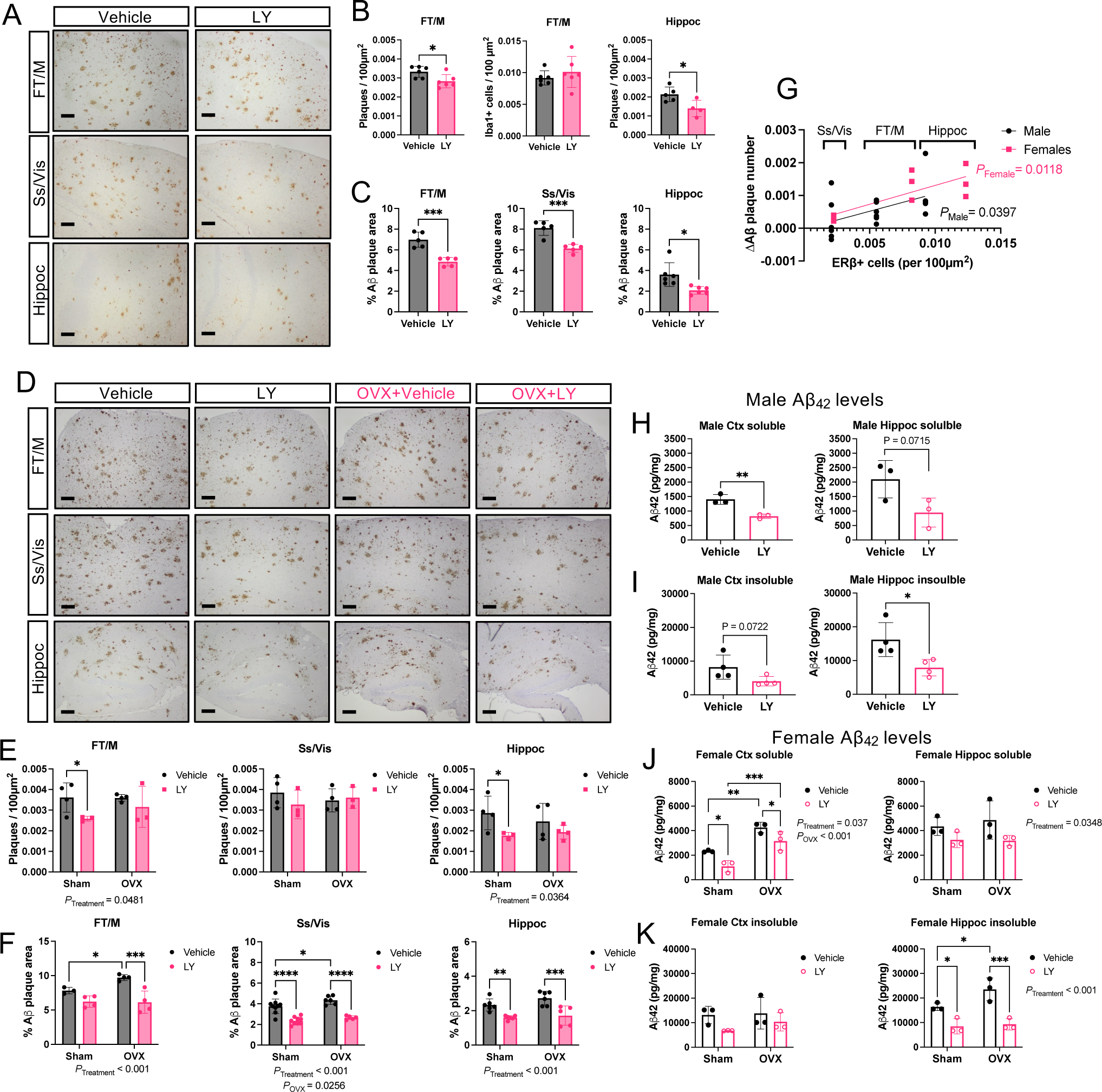
Less Aβ pathology in *App^NL-G-F^* male and female mice after ERβ activation. (A) Immunohistochemical representation of amyloid plaques in frontal and motor cortex (FT/M), somatosensory and visual cortex (Ss/Vis) and hippocampus (Hippoc) of male *App^NL-G-F^* mice after vehicle or LY treatment. (B) Quantification of number of plaques per 100 µm^2^ (n = 4-6) and (C) percent plaque area (n = 4-6) in male *App^NL-G-F^* mice. (D) Similar as in (A), immunohistochemical representation of amyloid plaques in different brain regions of female *App^NL-G-F^* mice after vehicle or LY treatment. (E) Quantification of number of plaques per 100 µm^2^ (n = 3-4) and (F) percent plaque area (n = 3-9) in female *App^NL-G-F^* mice. (G) Linear regression analysis comparing effect size from LY treatment (vehicle vs. LY) on number of Aβ plaques in relation to average number of ERβ positive cells per 100 µm^2^ in different brain regions of male and female mice (n = 3-6). (H) Soluble and (I) insoluble Aβ42 levels in male cortex (Ctx, left) and hippocampus (Hippoc, right) (n= 3-4). (J) Soluble and (K) insoluble Aβ_42_ levels in female cortex (Ctx, left) and hippocampus (Hippoc, right) (n = 3). * *P* < 0.05, ** *P* < 0.01, *** *P* < 0.001, **** *P* < 0.0001. Unpaired t-test was used for males and 2-way ANOVA for females followed by uncorrected Fisher’s LSD test for multiple comparisons. Overall significant main effects of treatment or OVX are indicated. Scale bars = 100 µm.

### 3.3. Effect of ERβ on APP processing

To explore if reduced Aβ_42_ levels in LY-treated mice was a consequence of lower APP levels or a shift from amyloidogenic β-secretase processing to non-amyloidogenic α-secretase processing, we analyzed the levels of full-length APP (FL-APP) and processed APP fragments. Western blot analysis revealed that LY-treatment of male mice had no effect on FL-APP levels in cortex nor in hippocampus (Fig. 3A-D). However, FL-APP was significantly increased in female cortex upon OVX and decreased upon LY-treatment of OVX females (Fig. 3A, B, C). LY treatment did not result in any difference in C-terminal fragment β-CTF levels relative to FL-APP (Fig. 3D), but β-CTF was increased upon OVX relative to β-actin levels (Fig. 3E), suggesting α-or β-secretase activities are not altered by ERβ activation (Fig. 3A, B, D). In addition, the expression of *App* or processing enzymes (*Bace1, Psen1,* and *Adam10)* were not altered by LY treatment or OVX (Supplemental Fig. 3C-J), further suggesting that ERβ does not modulate APP processing, although OVX increased APP protein levels in female cortex (Fig, 3A, C). Finally, we again observed lower total Aβ levels upon LY treatment in both male and female cortex and hippocampus and an interesting increase in cortex upon OVX (Fig. 3A, B, F). These data suggest that ERβ does not directly modulate APP processing but may rather be involved in clearance of Aβ.

**FIGURE 3.**
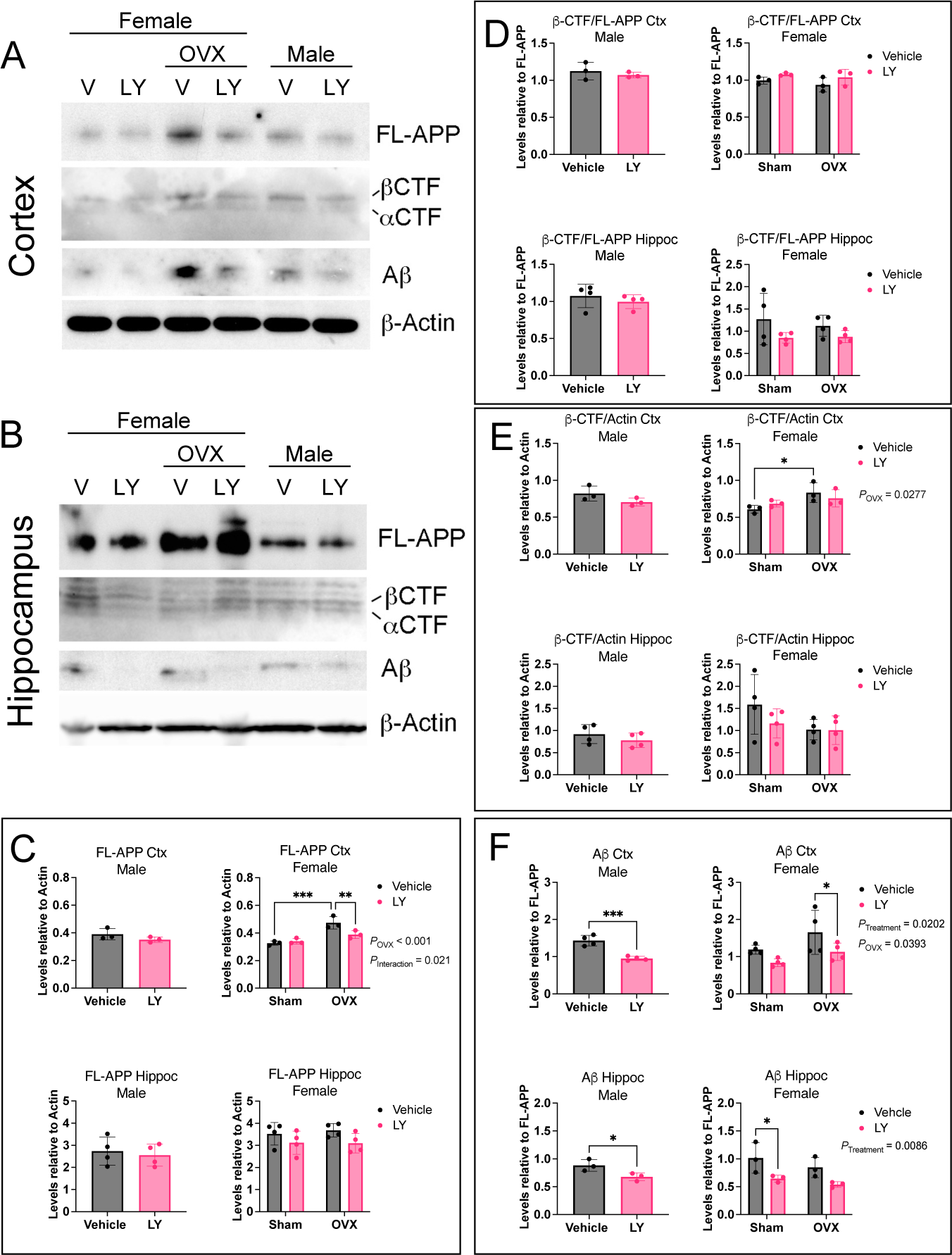
ERβ activation does not alter APP processing. Western blot analysis of full-length APP (FL-APP), β-CTF and α-CTF, Aβ peptide, and β-actin in (A) cortex and (B) hippocampus of female and male *App^NL-G-F^* mice after vehicle (V) or LY treatment, as well as after sham surgery or ovariectomy (OVX in females). Quantification of (C) FL-APP relative to β-actin, (D) β-CTF relative to FL-APP, (E) β-CTF relative to actin, and (F) Aβ relative to FL-APP in male (left), and female (right), cortex (Ctx) (top), and hippocampus (bottom) (n = 3-4). * *P* < 0.05, ** *P* < 0.01, *** *P* < 0.001. Statistical significance was determined using unpaired t-test for males and 2-way ANOVA for females followed by uncorrected Fisher’s LSD test for multiple comparisons. Overall significant main effects of treatment or OVX are indicated.

### 3.4. Effect of ERβ on glial cells in *App^NL-G-F^* mice

Astrocytes and microglia take active part in amyloid pathogenesis, including Aβ clearance (21). Therefore, we sought to investigate the impact of ERβ activation on astrocytic and microglial response. We could not detect any major effects on astrogliosis in male and female mice treated with LY or in OVX females (Supplemental Fig. 4A-D). Similarly, we did not see any difference in microglia numbers upon LY treatment in male *App^NL-G-F^* mice (Fig. 4A, B, Supplemental Fig. 5A). However, LY treatment significantly reduced number of activated CD68+ microglia, especially in the male hippocampus (Fig. 4A, C), and we also observed that LY treatment promoted microglia association to amyloid plaques in male frontal/motor cortex and in hippocampus (Fig. 4D). In female *App^NL-G-F^* mice, LY treatment had less effect on microglia (Fig. 4E-I, Supplemental Fig. 5B) compared to male mice, and we could detect a slight, but significant, effect of LY on lowering the number of activated CD68+ microglia in frontal/motor cortex and in hippocampus (Fig. 4H). We could not detect any significant effect of OVX on microglia numbers or activation. Interestingly, female mice had an overall increased number of CD68+ microglia, and overall increased number of plaque-associated microglia, but less response to LY (Supplemental Fig. 5C, D). Using a detection-panel of proinflammatory cytokines we observed significantly lower levels of IL-1b, CXCL1 (KC/GRO), and IL-12p70 in cortex and/or hippocampus of LY treated male mice (Fig. 5A-F). Of these, only IL-12p70 was significantly lower in female LY treated mice compared to vehicle treated mice and OVX sustained high levels of IL-12p70 in female hippocampus (Fig. 5E, F). Additionally, the level of the danger signal IL-10 was lower in LY-treated male hippocampus (Fig. 5G, H), which may suggest less hyperinflammation (22, 23) upon LY-treatment in males. Studying the expression of specific microglial markers, we could detect lower expression of *Cd68* and *Trem2* after LY treatment in female hippocampus (Fig 5I, J) and lower *Nos2* expression after LY treatment in male hippocampus (Fig. 5K). Interestingly, *Trem2* and *Cx3cr1* (that mediates microglial anti-inflammatory and pro-resolving responses) were significantly downregulated in female hippocampus upon OVX (Fig. 5J, L). No difference in these markers were observed in cortex (data not shown). Thus, ERβ may have a more direct and sex-specific effect on microglial function and thereby on Aβ scavenging.

**FIGURE 4.**
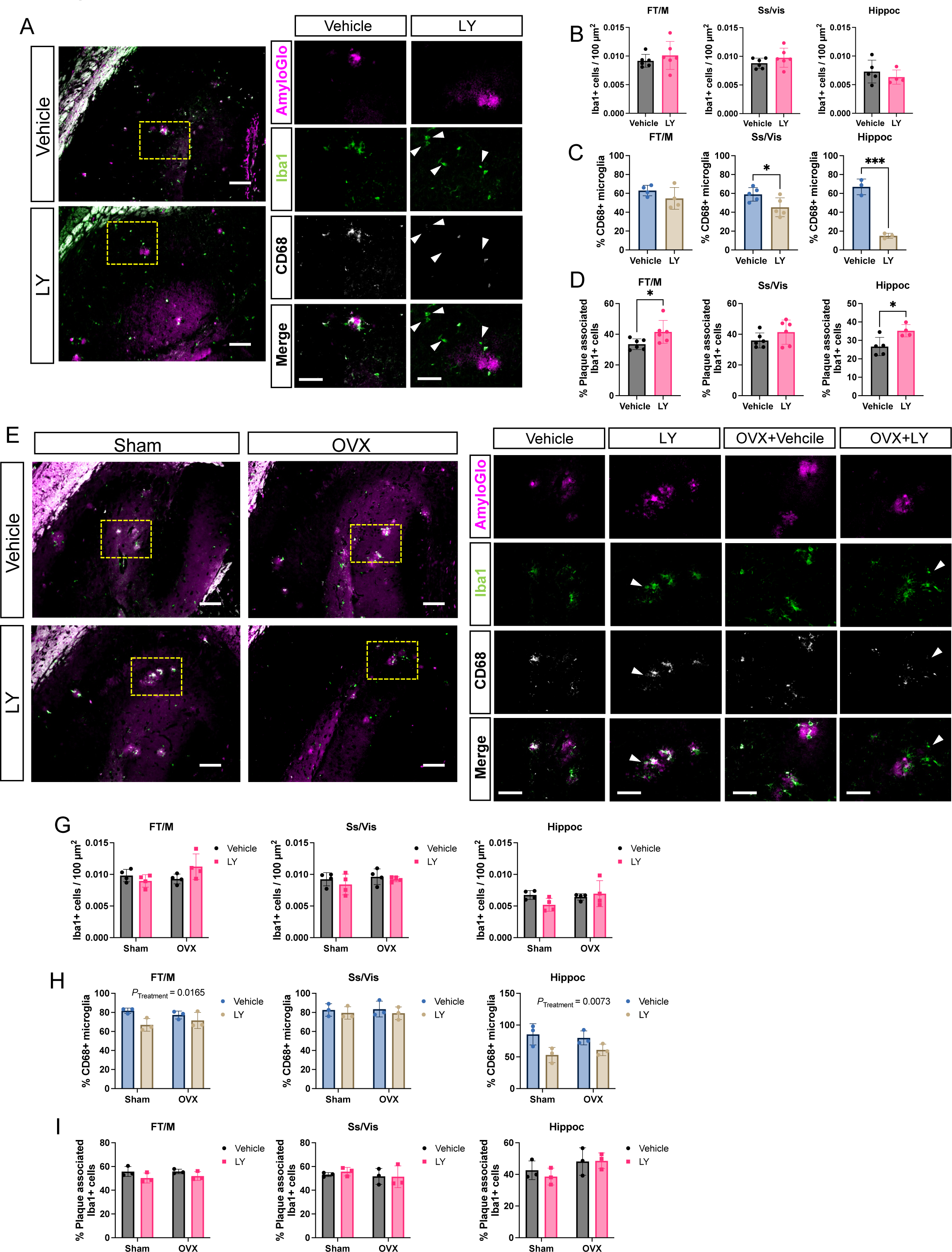
ERβ activation modulates microglia activation in a sex-specific manner in *App^NL-G-F^* mice. (A) Representative immunofluorescence images of male *App^NL-G-F^* hippocampus stained with the amyloid stain AmyloGlo (magenta), Iba1 (green), and CD68 (white) after vehicle or LY treatment. Yellow dotted area (left) indicates magnified region of interest (right). Arrowheads indicate microglia with lower CD68 levels. Scale bar 100 µm (left) and 50µm (right). Quantification in male *App^NL-G-F^* mice of (A) number of Iba1 cells per 100 µm^2^ (n = 4-5), (B) percent CD68+, Iba1+ double positive cells (n = 3-5), and (C) percent microglia within 20 µm radius of plaque edge (n = 4-6). Representative immunofluorescence images of female *App^NL-G-F^* hippocampus stained with AmyloGlo (magenta), Iba1 (green), and CD68 (white) after vehicle or LY treatment, as well as after sham surgery or ovariectomy (OVX). Yellow dotted area (left) indicates magnified region of interest (right). Arrowheads indicate microglia with lower CD68 levels. Scale bar 100 µm (left) and 50µm (right). Quantification in female *App^NL-G-F^* mice of (A) number of Iba1 cells per 100 µm^2^ (n = 4), (B) percent CD68+, Iba1+ double positive cells (n = 3), and (C) percent plaque-associated microglia (n = 3). * *P* < 0.05, *** *P* < 0.001. Unpaired t-test was used for males and 2-way ANOVA for females followed by uncorrected Fisher’s LSD test for multiple comparisons. Overall significant main effects of treatment are indicated.

**FIGURE 5.**
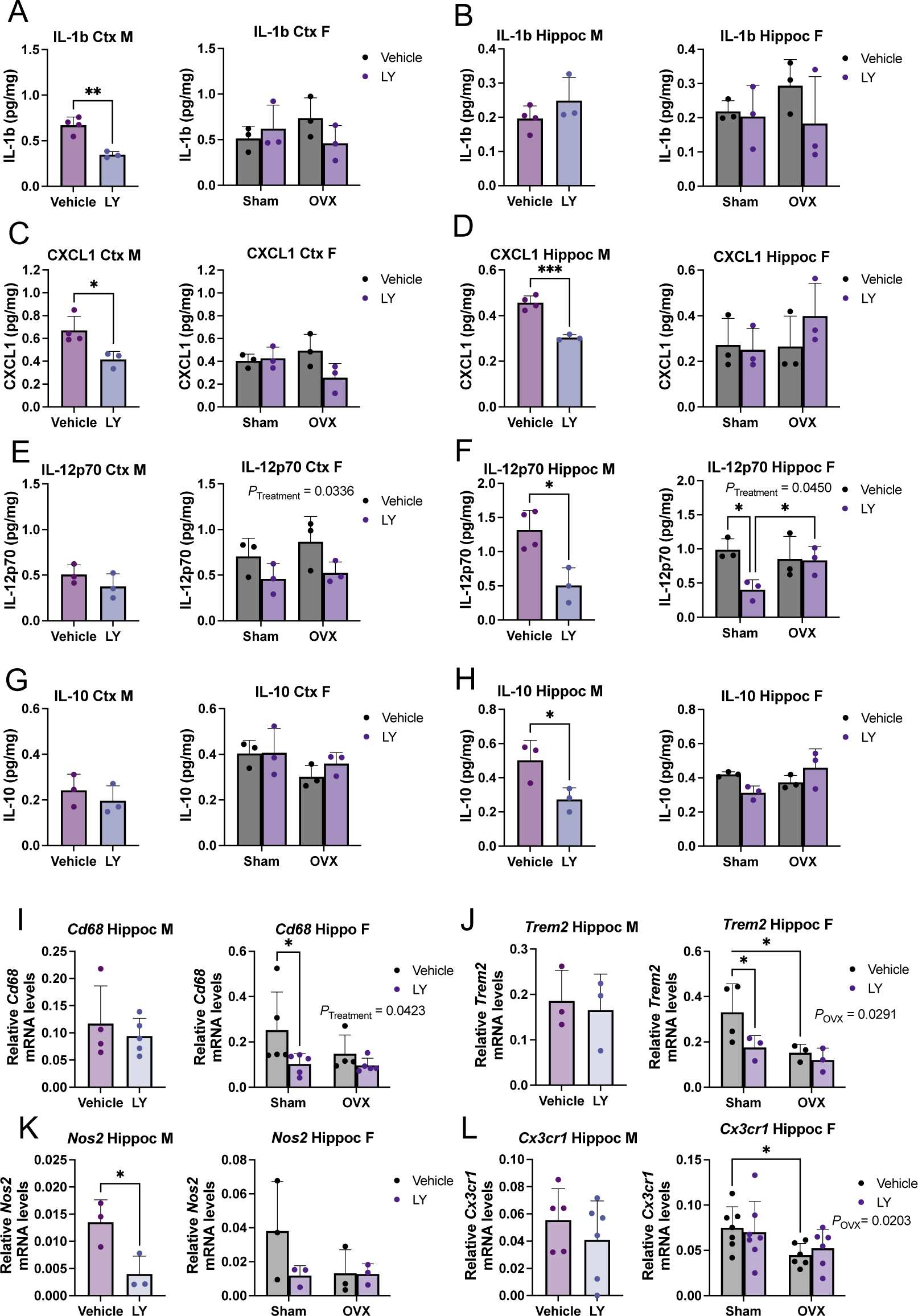
Microglial and proinflammatory markers are altered upon ERβ activation in a sex-specific manner. Multiplex ELISA analysis of the proinflammatory markers IL-1b, CXCL1 (KC/GRO), and IL-12p70, as well as IL-10, in male (left) and female (right) *APP^NL-G-F^* cortex (Ctx) (A, C, E, G) and hippocampus (Hippo) (B, D, F, H), after vehicle or LY treatment, as well as after sham surgery or ovariectomy (OVX in females) (n = 3-4). Expression of microglial and inflammatory markers (I) *Cd68*, (J) *Nos2,* (K) *Trem2*, and (L) *Cx3cr1* relative to housekeeping gene *Rplp0* expression in male (left) and female (right) *App^NL-G-F^* hippocampus after vehicle or LY treatment, as well as after sham surgery or OVX in females (n = 3-7). * *P* < 0.05, ** *P* < 0.01, *** *P* < 0.001. Unpaired t-test was used for males and 2-way ANOVA for females followed by uncorrected Fisher’s LSD test for multiple comparisons. Overall significant main effects of treatment or OVX are indicated.

### 3.5. Loss of ERβ modulates microglial function in a sex-specific manner

To study any direct effects of ERβ on microglial function, we first analyzed co-expression of ERβ with Iba1+ microglia in WT and *Esr2*-KO mice. Although some microglia showed ERβ positive staining, most microglia were ERβ negative (Fig. 6A). In *App^NL-G-F^* mice, plaque associated microglia were both ERβ positive and negative and LY treatment had no overall effect on ERβ-Iba1 co-expression in any brain region examined (Fig. 6B, Supplemental Fig. 6A, B). However, there were slightly but significantly more ERβ+ microglia in male compared to female *App^NL-G-F^* brains (Supplemental Fig. 6C). We next isolated primary microglia from WT and *Esr2*-KO adult male and female mice to study any effect of ERβ on microglia function. Microglia originating from both male and female WT mice express ERα (*Esr1*) and ERβ (*Esr2*) mRNA, of which *Esr1* levels were ca 10-fold higher than *Esr2* and were also higher in female compared to male primary microglia, as well as higher in *Esr2*-KO microglia (Supplemental Fig. 6D-F). The primary microglial cultures were treated with or without LY 24h prior to, and during, 24h M1 (pro-inflammatory), M2 (anti-inflammatory) or control stimulation, followed by fluorescent Aβ incubation. Loss of ERβ decreased overall Aβ phagocytosis in male microglia, while LY treatment had minor effects (Fig. 6C). The effect of ERβ loss was not as evident on phagocytosis in female microglia although there was an interesting LY-mediated decrease of M1 activated phagocytosis even in *Esr2*-KO microglia (Fig 6D), which could be an off-target effect of ERα activation since *Esr1* levels were increased in female and *Esr2*-KO microglia (Supplemental Fig 6E, F). Microglia migration towards CX3CL1 (a ligand for CX3CR1) was not affected by LY treatment or ERβ loss in male microglia (Fig. 6E). Female microglia migrated less upon ERβ loss in the control condition and no clear effect of LY could be observed (Fig. 6F). Comparing male and female WT microglia we could observe that male microglia were overall more phagocytic but less migratory than female microglia (Supplemental Fig. 6G, H). Next, we explored mRNA expression of key microglial markers. *Nos2* expression was highly elevated in the M1 condition where LY treatment lowered its expression in male but not female microglia (Fig. 6G, H), however without any significant effect of ERβ loss. Markers expressed under control and M2 conditions included *Trem2* and *Cx3Cr1*. *Trem2* was overall increased in female but not male microglia upon ERβ loss (Fig. 6I, J), while ERβ loss decreased *Cx3cr1* (that is involved in microglial activation and phagocytosis) in male microglia at least under control conditions (Fig. 6K, L), which is in line with reduced phagocytosis in male *Esr2*-KO microglia. In contrast, *P2ry12*, a factor needed for microglial chemotaxis, was overall slightly lower in female microglia upon ERβ loss (but not in male microglia) (Fig. 6M, N), which could reflect the lower migration of female *Esr2*-KO microglia. Although the typical M2 marker *Arg1* was higher expressed upon M2 treatment (Fig 6O, P), it appears that the primary microglia had a partial M2 phenotype already under the control condition, also reflected by increased *Trem2* and *Cx3cr1* expression (Fig. 6I-L). Finally, the more general microglia activation marker *Cd68* was overall higher expressed upon ERβ loss in male microglia (Fig. 6Q, R). Although we did not see any effect of LY on *Cd68* expression, this data combined with the lower levels of Cd68+ microglia in LY-treated *App^NL-G-F^* mice points to a role of ERβ in microglia activation, especially in male mice. Combined, our data show sex-dimorphic effects from ERβ loss in primary microglia, but these effects are overall mild suggesting that other cell types than microglia are also involved in ERβ’s neuroprotective functions.

**FIGURE 6.**
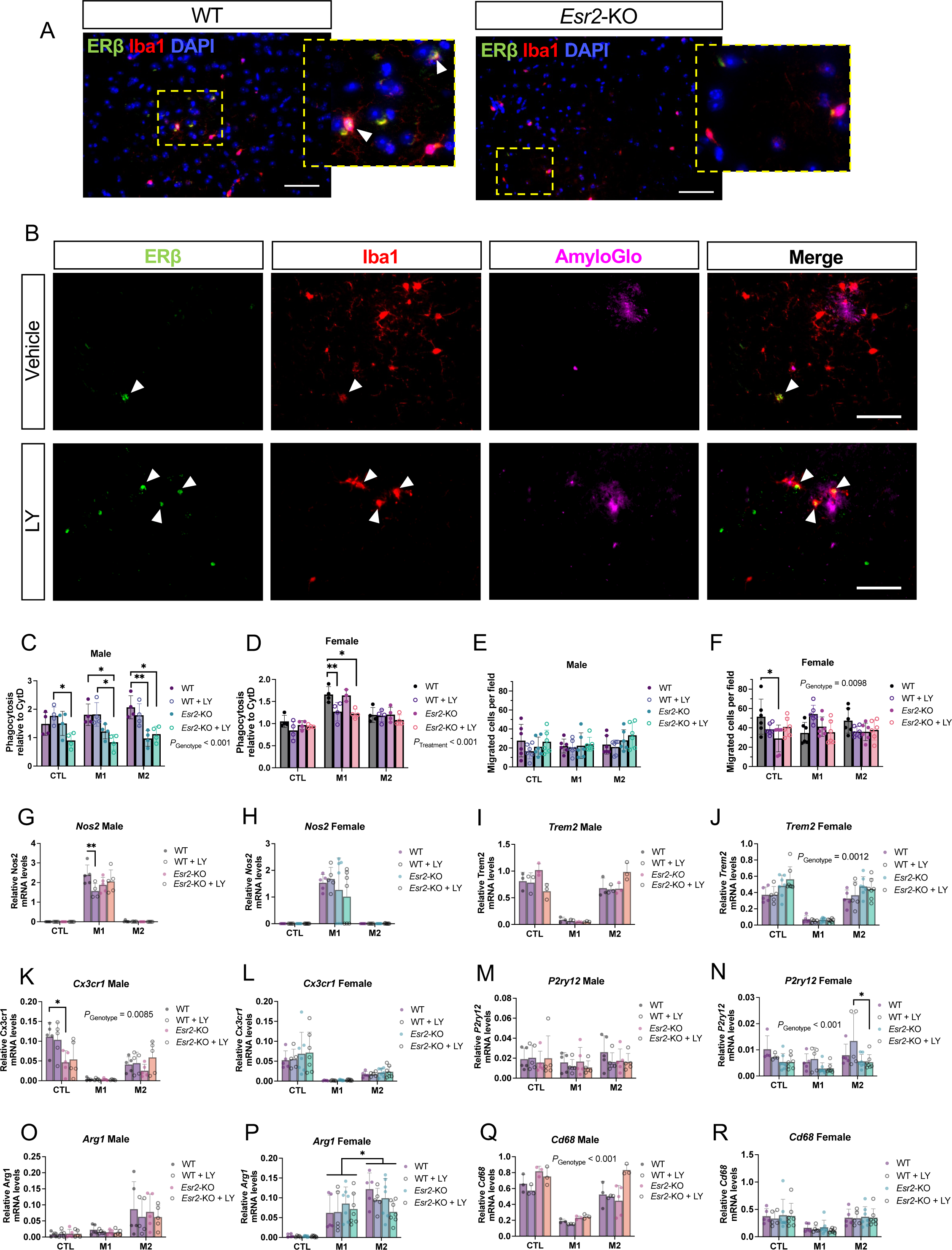
Loss of ERβ modulates sex-specific effects in primary mouse microglia. (A) Representative immunofluorescence of ERβ and Iba1 co-staining (arrowheads) in WT (left) and *Esr2*-KO (right) male cortex (dotted rectangle: magnified area), and (B) in male *App^NL-G-F^* cortex upon vehicle or LY treatment (scale bars = 50 µm). Phagocytosis of Aβ (relative to Cytochalasin D treatment) of primary microglia originating from male (C) and female (D) WT and *Esr2*-KO mice. The microglia were treated with or without 10 nM LY 24h prior to and during 24h M1, M2, or no (CTL) activation stimulation (n = 3-4). Transwell migration assay of primary microglia originating from male (E) and female (F) WT and *Esr2*-KO mice treated as indicated above (n = 6). Expression of microglial markers *Nos2*, *Trem2*, *Cx3cr1*, *P2ry12*, *Arg1*, and *Cd68* (G-R) relative to housekeeping gene *Rplp0* expression in primary microglia originating from male (G, I, K, M, O, Q) and female (H, J, L, N, P, R) WT and *Esr2*-KO mice treated as indicated above (n = 3-7). * *P* < 0.05, ** *P* < 0.01. Unpaired t-test was used for male microglia and 3-way ANOVA or mixed-effects analysis for female microglia followed by Tukey’s or Dunnett’s multiple comparison test. Overall significant main effects of LY treatment or genotype are indicated.

## 4. DISCUSSION

Estrogen signaling has been ascribed neuroprotective properties in AD (1-3). In this study we explored specifically the role of ERβ in mediating neuroprotection in the *App^NL-G-F^* mouse model of AD, a model that unlike previous AD models circumvents the artefacts of APP overexpression, making it more clinically relevant (13). We show that selective ERβ activation with LY500307 (LY) protects against amyloid pathology and memory deficits in *App^NL-G-F^* mice. We also show that this neuroprotection is different between males and females, likely involving different cell types, and that ovariectomy (OVX) increases Aβ_42_ levels but has otherwise a limited effect on overall pathology in *App^NL-G-F^* mice.

Despite previous problems with ERβ antibody specificities, it is now clear that both ERα and ERβ are expressed in the cortex and hippocampus but in a scattered manner and at relatively low levels (1). E2 has been ascribed general neuroprotective effects by protecting against apoptosis (24), and sustaining mitochondrial health and thereby regulating oxidative stress (2, 25, 26). E2 also promotes neurogenesis and synaptic plasticity upstream of BDNF (27, 28) and WNT signaling (29, 30). Although these pathways likely also contribute to estrogenic neuroprotection in AD, very few studies exist on the role of ERs in animal AD models, and even fewer address possible sex differences in estrogenic neuroprotection. However, it has been shown that ERα activation protects against memory deficits in female APP/PSEN1 transgenic mice(31), and reduces Aβ accumulation in female 3xTg-AD transgenic mice (10). Similarly, ERβ activation using dietary phytoestrogens (with various specificity to ERβ) lowers Aβ deposition and ameliorates cognitive deficits in female APP/PSEN1 mice (9, 32, 33) and in female (11) and male (34) 3xTg mice, which in part could be attributed to modulated BDNF and WNT signaling, and enhanced microglial phagocytosis (11, 33). Importantly, a direct comparison between ERβ activation in male and female AD models have until now been missing.

Early loss of circulating estrogen and progesterone such as in early menopause or bilateral oophorectomy may be a risk factor for AD (4-6), and E2 supplementation could protect against this risk. Human data on such protective associations are limited and controversial (35). However, animal studies using 3xTg-AD mice demonstrate that gonadoectomy leads to increased Aβ accumulation and cognitive impairment, while estrogenic supplementation protects against these deficits (10, 34, 36-38). Interestingly, similar protection was not seen in ovariectomized APP/PSEN1 mice (39), suggesting that inherent model characteristics may modulate estrogenic neuroprotection.

In the present study we used the *App^NL-G-F^* model that express human Aβ with knock-in of *NL-G-F* mutations into the endogenous mouse *App* gene (13), circumventing *APP* overexpression artefacts of earlier transgenic AD models. By selectively activating ERβ in this model, we not only confirm previous studies in older AD transgenic models on ERβ’s protective effects (9, 11, 32, 33), we also identify important new sex-differences in ERβ mediated protection. However, in contrast to 3xTG AD models, OVX did not yield major effects on AD pathology or memory function in our study. Interestingly, the only clear effects of OVX were increased soluble and insoluble Aβ_42_ levels in cortex and hippocampus, respectively (Fig. 2J, K), which could be related to higher FL-APP levels in OVX mice (Fig. 3A, E) (with an interesting interaction between OVX and LY on cortical FL-APP levels (Fig. 3C)), and decreased *Trem2* expression in hippocampus, a microglial protein participating in Aβ clearance (Fig. 5J).

Our study also suggests that ERβ works differently with different effect sizes in cortex and hippocampus. Although ERβ mRNA expression levels were similar between cortex and hippocampus, a more detailed brain region analysis showed that the number of ERβ+ cells was highest in frontal and primary motor cortex as well as in hippocampus (Supplemental Fig. 1A, B), which overlapped with the brain regions with largest effects of LY on Aβ plaque numbers (Fig. 2G). Furthermore, although the effect of OVX appeared to be different between cortex and hippocampus (Fig. 2F, J, K, 3F), ERβ activation lowered Aβ levels in both sham and OVX mice but had less clear role on neuroinflammation. Thus, more studies are needed to explain the impact of OVX in different brain cells and brain regions and its interaction with specific estrogen receptors. It is known that OVX modulates glucose metabolism differently in different brain regions (40), and that E2 can be de-novo synthesized in different brain regions (including hippocampus and cortex) and in different cell types (41, 42). Similarly, it is likely that ERβ mediates brain region-specific functions through interactions with different cell type-specific factors. For example, ERβ (but not ERα) can regulate BDNF signaling in the female rodent brain in a region-specific manner (28). In addition, since OVX does not only affect ERβ signaling, but all ER signaling, direct relationships between LY treatments and OVX cannot be expected.

Our data showed no effect of ERβ on APP processing. Since a previous study demonstrated a role for ERβ in microglia activation (43), we turned to examine if Aβ scavenging could be affected by ERβ. ERβ activation markedly reduced microglia activation in both male and female mice, with the strongest effect in the male hippocampus, which was concomitant with decreased levels of proinflammatory markers (Fig. 5). This may mean that ERβ activation leads to less amyloidosis and therefore less neuroinflammation. However, ERβ activation also increased number of plaque-associated microglia in at least hippocampus (Fig. 4), which argues for a more direct effect. Although, this effect was largely sex-specific, with the strongest effect in male hippocampus, an interesting effect in females of OVX on lowering expression of two microglial markers involved in chemotaxis and phagocytosis, *Cx3cr1* and *Trem2* (Fig. 5J, L), suggest that circulating estrogen, acting likely via both ERβ and ERα, may be needed to sustain microglia function, which is in line with that a minority of microglia were ERβ positive in our study (Fig. 6A, B, Supplemental Fig. 6A-C).

To anyway elaborate on a possible direct role of ERβ on microglia, we used primary microglia from *Esr2*-KO and WT male and female mice. Again, we could observe sex differences in how ERβ modulates the function of microglia where loss of ERβ resulted in decreased phagocytosis in male-originating microglia, but no effect on migration, while ERβ loss in female-originating microglia resulted in slightly decreased migration but no effect on phagocytosis (Fig 6A-D). In these primary microglia, the difference between male and female originating microglia are the sex chromosomes, suggesting that ERβ possibly interact with genes on the Y-chromosome to elicit the sex-specific effect of ERβ loss. In addition, effect of LY was minimal on the primary microglia (except lowering *Nos2* expression in male microglia), implying that ERβ largely works ligand-independent in at least primary microglia setting. However, LY lowered phagocytosis in M1 activated female microglia, even in *Esr2*-KO microglia, suggesting that LY may target ERα cell-type specifically when ERβ levels are low (Fig 6B, Supplemental Fig. 6A-C), which is an important finding when considering possible ERβ targeting therapies. A drawback with our primary microglia culture is that we, despite optimization, acquire a basal M2-like phenotype even in the control condition. This could mask effects of LY or ERβ loss especially upon M1 activation. A further limitation of the microglia culture is that we had to culture the cells in presence of normal FBS since microglia did not tolerate steroid-stripped FBS, which could also have masked effects of LY. Combined with our data that only a fraction of microglia express ERβ in the *App^NL-G-F^* mouse brain, we hypothesize that ERβ can exert sex-specific neuroprotective effects via microglia but that other cell-types are likely also involved. Thus, our study emphasizes the sex differences in ERβ’s neuroprotection; in male mice this neuroprotection can be mediated through microglia, while in female mice other non-inflammatory processes downstream of ERβ activation appear to play a larger role. Autophagy may be such process as suggested by Wei and coworkers (12).

In conclusion, our study provides the first direct comparison of ERβ’s sex-specific neuroprotective effects in an AD model. We show that this neuroprotection is not directly associated with altered APP processing, but rather in part mediated via microglia in a sex-specific manner, and that ovariectomy can increase Aβ levels, but had otherwise limited overall effects on AD pathology. Our research adds to the molecular understanding to the sex-differences in AD and warrants further studies on brain cell-specific effects of ERβ in male and female AD models and in human AD patients.

## ACKNOWLEDGEMENTS

We thank Wanfu Wu, Julian Jung, and Johanna Wanngren for excellent technical assistance. We also thank Takashi Saito and Takaomi Saido at RIKEN Center for Brain Science for providing *App* knock-in mice, the Animal Behavior Core Facility (ABCF) of Karolinska Institutet where the behavioral studies were performed, and KI Stem Cell Organoid Facility (KISCO) for help with primary microglia experiments.

## DATA AVAILABILITY

Any information required to analyse the data in this paper is available from authors upon reasonable request.

