## Supplemental material for "ERβ mediates sex-specific protection in the *App-NL-G-F* mouse model of Alzheimer’s disease"

**SUPPLEMENTAL TABLE 1.**

Antibody list.

**SUPPLEMENTAL TABLE 2.**

Real-time qPCR primer and assay list.

**SUPPLEMENTAL FIGURE 1.**

ERα and ERβ expression in the mouse cortex and hippocampus

**SUPPLEMENATL FIGURE 2.**

Effect of LY treatment on $A\beta_{40}$ levels in $App^{NL\text{-}G\text{-}F}$ mice.

**SUPPLEMENTAL FIGURE 3.**

APP processing and expression of APP processing enzymes in $App^{NL\text{-}G\text{-}F}$ mice.

**SUPPLEMENTAL FIGURE 4.**

Effect of ERβ activation on astrocyte numbers in $App^{NL\text{-}G\text{-}F}$ mice.

**SUPPLEMENTAL FIGURE 5.**

ERβ activation modulates microglia in $APP^{NL\text{-}G\text{-}F}$ mice.

**SUPPLEMENTAL FIGURE 6.**

ER expression and effect of ERβ knockout (*Esr2*-KO) in male and female mouse microglia.

**Supplemental Table 1.** Antibody list

| Antigen | Clone / catalogue number | Supplier | Dilution | Application |
|---|---|---|---|---|
| ERβ | PP-PPZ506-00 | R&D Systems | 1:5000 | IHC, IF |
| Aβ 1-42 | 18582 | IBL | 1:2000 | IHC |
| APP (N-terminal) | MAB348 (22C11) | Millipore | 1:1000 | Western Blot |
| APP (Aβ) | 6E10 | BioLegend | 1:800 | Western Blot |
| APP (C-terminal) | A8717 | Sigma-Aldrich | 1:1000 | Western Blot |
| β-Actin | A2228 | Sigma-Aldrich | 1:10 000 | Western Blot |
| GFAP | MAB3402X | Chemicon | 1:200 | IF |
| CD68 | ab283654 | Abcam | 1:200 | IF |
| Iba1 | ab178846 | Abcam | 1:400 | IF |

IHC: Immunohistochemistry
IF: Immunofluorescence

**Supplemental Table 2.** Real-time qPCR primer and assay list

TaqMan assay sets (ThermoFisher)

| | |
|---|---|
| *Rplp0* | AR3227N |
| *Esr1* | Mm00433149_m1 |
| *Esr2* | Mm00599821_m1 |

qPCR primers

| Gene | Fw (5'-3') | Rev (5'-3') |
|---|---|---|
| *App* | TCCGAGAGGTGTGCTCTGAA | CCACATCCGCCGTAAAAGAATG |
| *Psen1* | GGTGGCTGTTTTATGTCCCAA | CAACCACACCATTGTTGAGGA |
| *Bace1* | CAGTGGGACCACCAACCTTC | GCTGCCTTGATGGACTTGAC |
| *Adam10* | ATGGTGTTGCCGACAGTGTTA | GTTTGGCACGCTGGTGTTTTT |
| *Cd68* | TGTCTGATCTTGCTAGGACCG | GAGAGTAACGGCCTTTTTGTGA |
| *Cx3cr1* | CGGCCATCTTAGTGGCGTC | GGATGTTGACTTCCGAGTTGC |
| *Il4* | GGTCTCAACCCCCAGCTAGT | GCCGATGATCTCTCTCAAGTGAT |
| *Nos2* | GTTCTCAGCCCAACAATACAAGA | GTGGACGGGTCGATGTCAC |
| *Trem2* | CTGGAACCGTCACCATCACTC | CGAAACTCGATGACTCCTCGG |
| *P2ry12* | ATGGATATGCCTGGTGTCAACA | AGCAATGGGAAGAGAACCTGG |
| *Arg1* | CTCCAAGCCAAAGTCCTTAGAG | AGGAGCTGTCATTAGGGACATC |
| *Rplp0* | ACCTCCTTCTTCCAGGCTTT | CCCACCTTGTCTCCAGTCTTT |

### Supplemental Figure 1

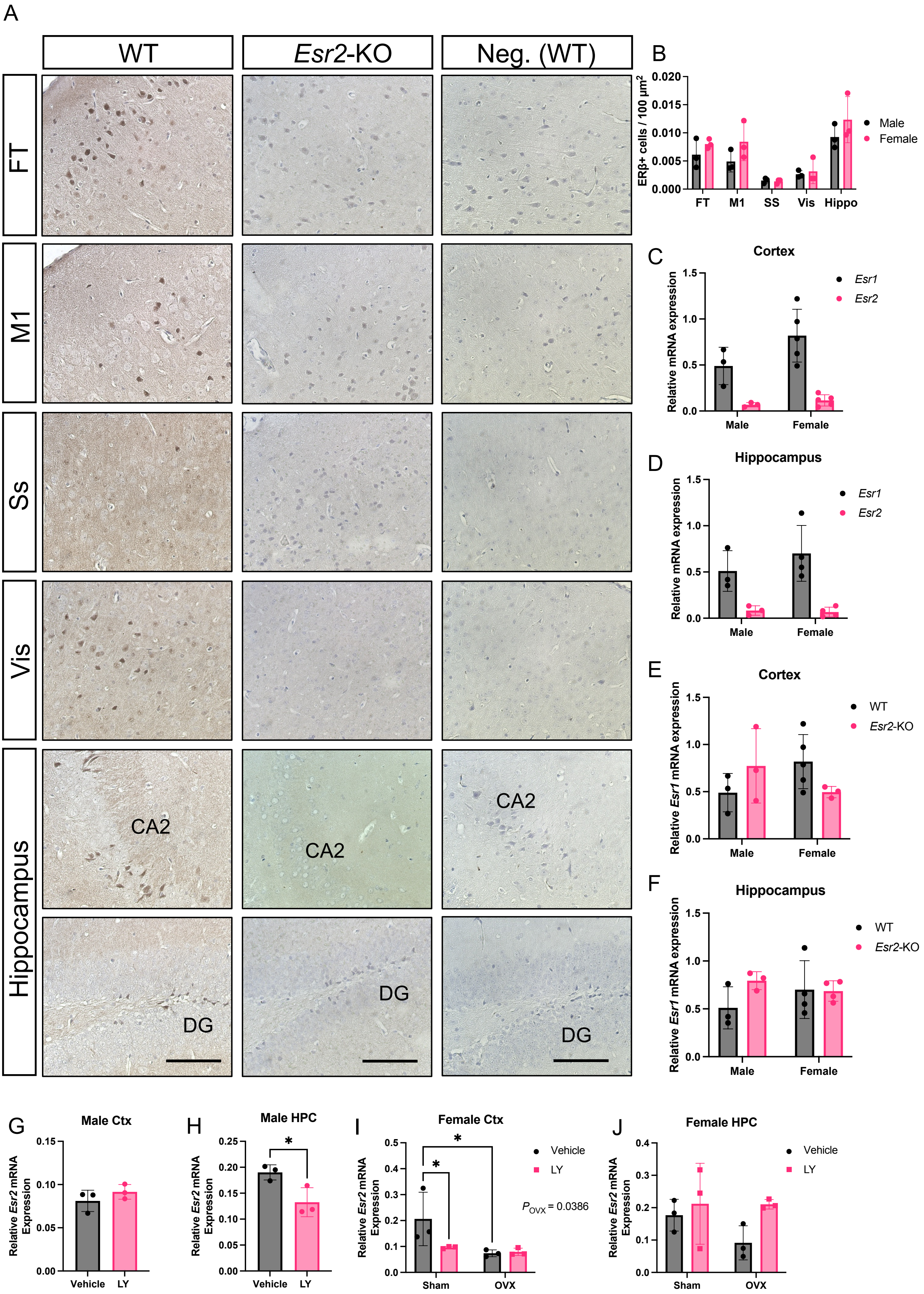

**SUPPLEMENTAL FIGURE 1. ERα and ERβ expression in the mouse cortex and hippocampus.** (A) Representative immunohistochemical images of ERβ in frontal (FT), primary motor (M1), somatosensory (Ss), and visual cortex (Vis), as well as in hippocampus of WT and ERβ knockout (*Esr2*-KO) mice. Negative control is without primary antibody. CA2 and dentate gyrus (DG) regions are indicated (scale bar = 100 µm). (B) Number of ERβ+ cells in male and female WT mice per 100 $\mu m^2$ of respective brain area. In hippocampus (Hippoc) the area of DG and CA1-3 were quantified for ERβ+ cells (n=3). *Esr1* (ERα) and *Esr2* (ERβ) mRNA expression in (C) cortex and (D) hippocampus of male and female mice. *Esr1* mRNA expression in (E) cortex and (F) hippocampus of WT and *Esr2*-KO male and female mice. *Esr2* mRNA expression after vehicle or LY500307 (LY) treatment in (G) male cortex, (H) male hippocampus (HPC), (I) female cortex, and (J) female hippocampus of $App^{NL\text{-}G\text{-}F}$ mice. All mRNA expression is shown relative to *Rplp0* reference gene (n=3). Statistical significance was determined using unpaired t-test was used (in G, H) and 2-way ANOVA followed by Tukey's multiple comparisons test (in C-F) or uncorrected Fisher's LSD test (in I, J). Overall significant main effect of OVX is indicated. * $P < 0.05$.

### Supplemental Figure 2

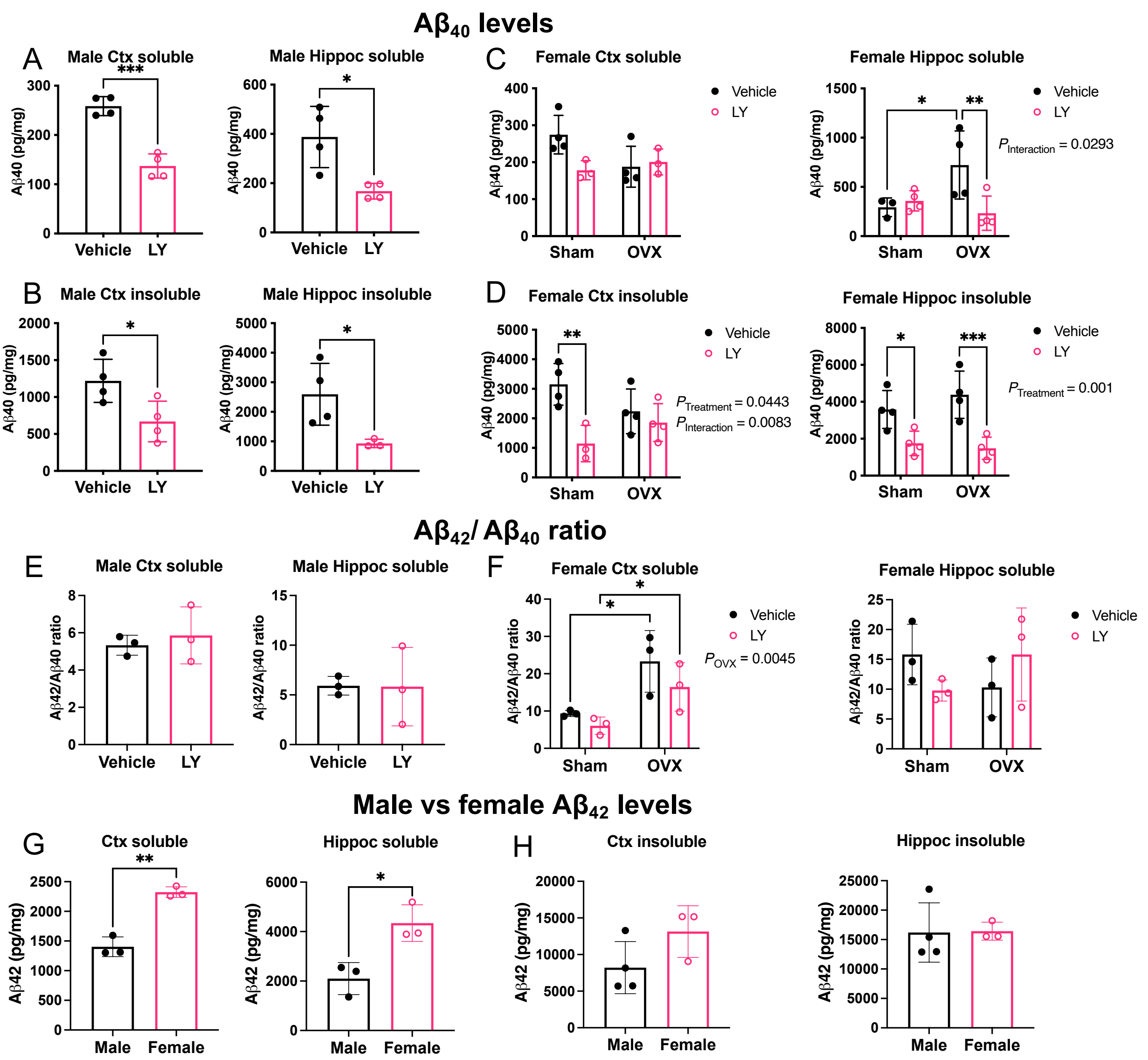


**SUPPLEMENATL FIGURE 2. Effect of LY treatment on A$\beta_{40}$ levels in *App*$^{NL\text{-}G\text{-}F}$ mice.** (A) Soluble and (B) insoluble A$\beta_{40}$ levels in male cortex (Ctx, left) and hippocampus (Hippoc, right) (n= 3). (C) Soluble and (D) insoluble A$\beta_{40}$ levels in female cortex (left) and hippocampus (right) (n = 3). Ratio of soluble A$\beta_{42}$/A$\beta_{40}$ in (E) male and (F) female cortex (Ctx, left) and hippocampus (Hippoc, right) (n= 3). Females were sham operated or ovariectomized (OVX). Comparison between male and female (G) soluble and (H) insoluble A$\beta_{42}$ levels in cortex (left) and hippocampus (right) (n = 3-4). * $P < 0.05$, ** $P < 0.01$. Unpaired t-test was used for males (and comparisons between male and females) and 2-way ANOVA for females followed by uncorrected Fisher's LSD test. Overall significant effects of treatment and interactions between treatment and OVX are indicated.

### Supplemental Figure 3

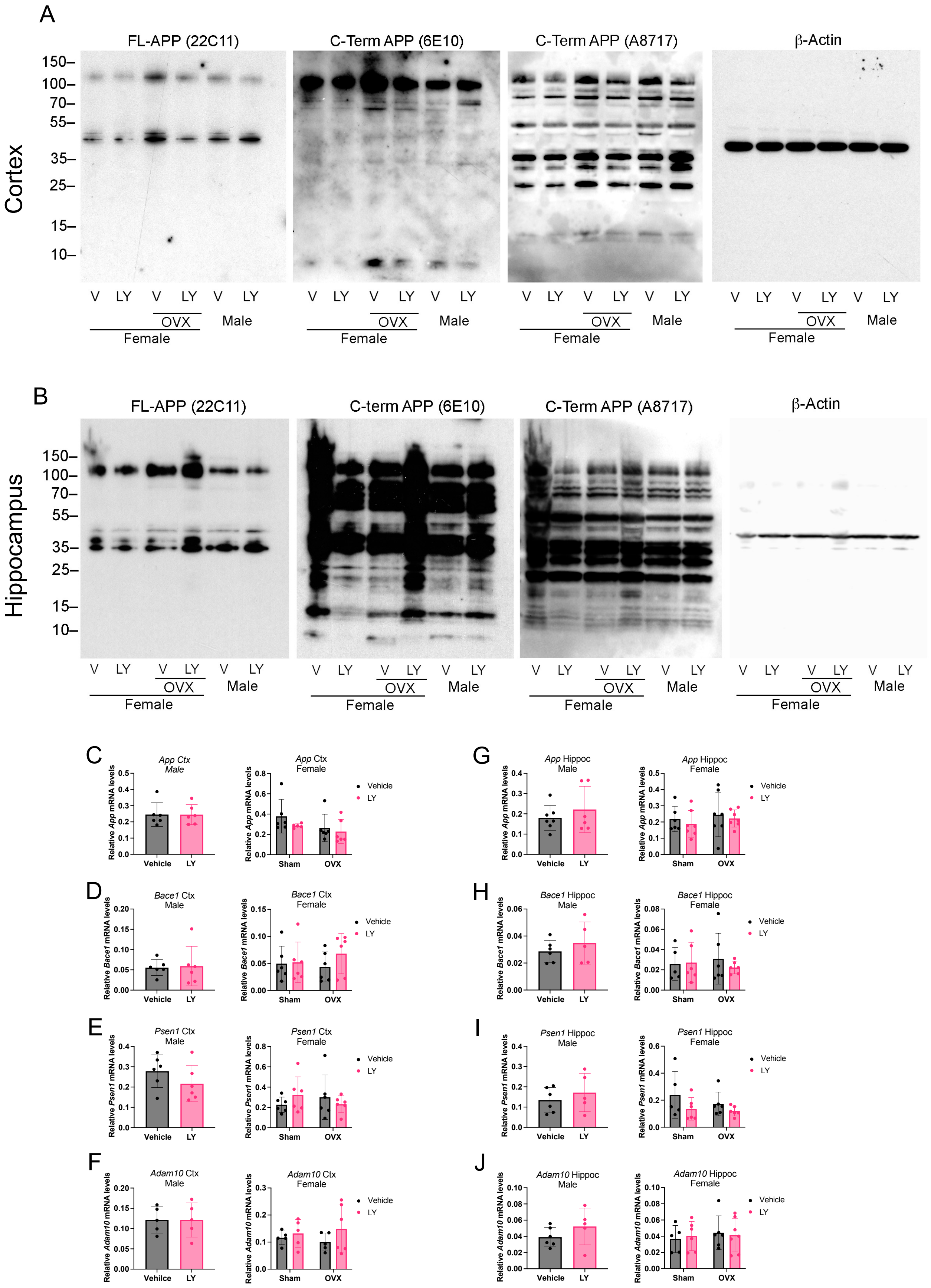

**SUPPLEMENTAL FIGURE 3. APP processing and expression of APP processing enzymes in *App*$^{NL\text{-}G\text{-}F}$ mice**. (A-B) The full-length blots for Figure 3. mRNA expression relative to *Rplp0* reference gene of *App*, *Bace1*, *Psen1*, and *Adam10* in cortex (C-F) and hippocampus (G-J) in male (left) and female (right) *App*$^{NL\text{-}G\text{-}F}$ mice treated vehicle or LY. Females were sham operated or ovariectomized (OVX). Unpaired t-test was used for males and 2-way ANOVA for females followed by uncorrected Fisher's LSD test.

### Supplemental Figure 4

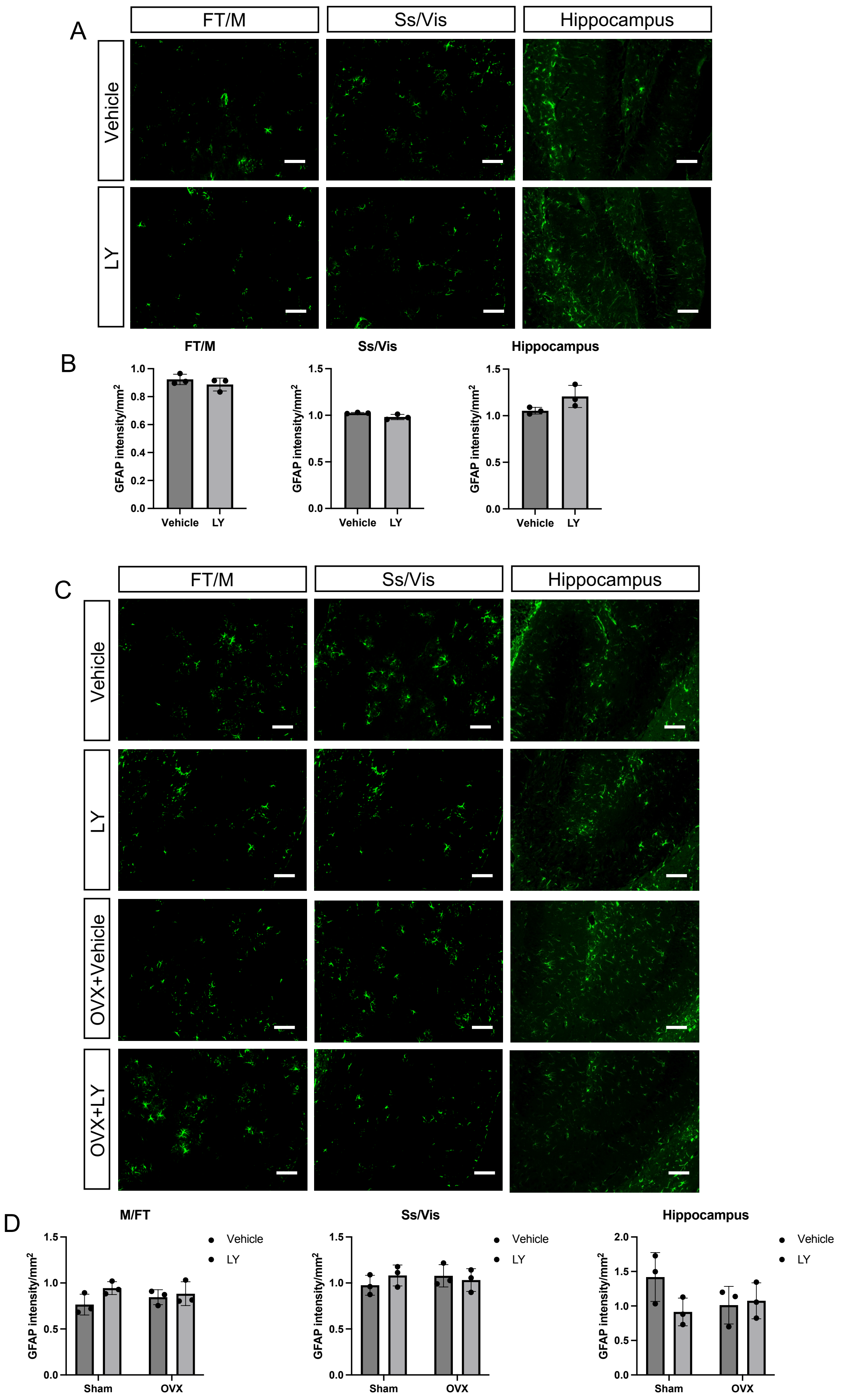

**SUPPLEMENTAL FIGURE 4. Effect of ERβ activation on astrocyte numbers in *App*$^{NL\text{-}G\text{-}F}$ mice.** Representative immunofluorescence images and quantification of GFAP (green) frontal/motor cortex (FT/M), somatosensory/visual cortex (Ss/Vis), and in hippocampus in (A-B) male and (C-D) female *App*$^{NL\text{-}G\text{-}F}$ mice after vehicle or LY treatment (scale bar = 100 µm, n = 3). Females were sham operated or ovariectomized (OVX). Unpaired t-test was used for males and 2-way ANOVA for females followed by uncorrected Fisher's LSD test.

### Supplemental Figure 5

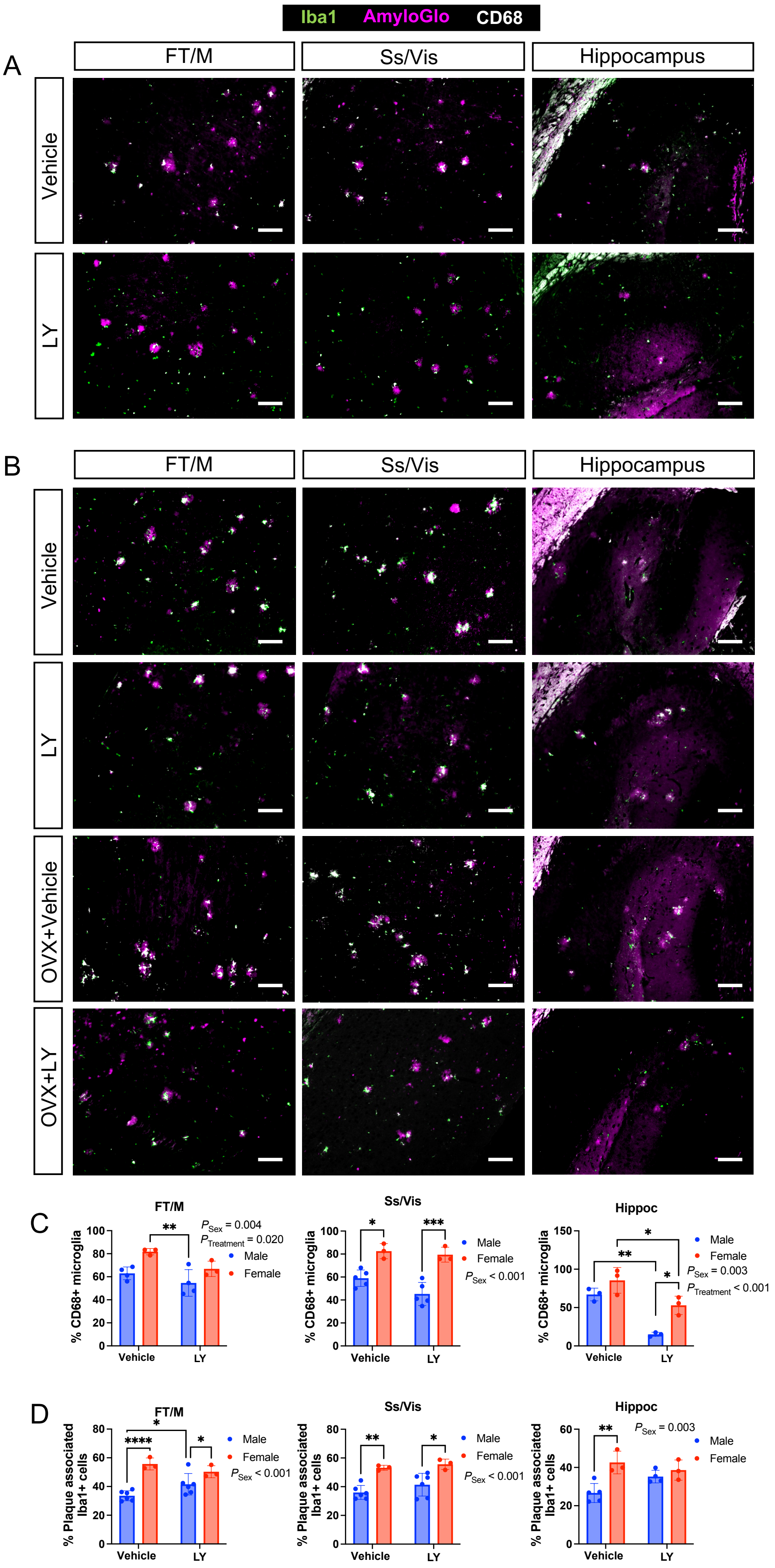

**SUPPLEMENTAL FIGURE 5. ERβ activation modulates microglia in *APP*$^{NL\text{-}G\text{-}F}$ mice.** Representative immunofluorescence images related to Figure 4 of frontal/motor cortex (FT/M), somatosensory/visual cortex (Ss/Vis) and hippocampus in (A) male and (B) female *APP*$^{NLGF}$ mice after vehicle or LY treatment (scale bar = 100 µm). Females were sham operated or ovariectomized (OVX). (C) Quantification of CD68+ microglia (n = 3-5) and (D) % plaque associated microglia expression (n = 3 – 6) in FT/M, Ss/Vis and hippocampus of male and female *App*$^{NL\text{-}G\text{-}F}$ mice. Statistical significance was determined using 2-way ANOVA followed by Tukey's multiple comparisons test. Overall significant effects of sex and treatment are indicated. * $P < 0.05$, ** $P < 0.01$, *** $P < 0.001$

### Supplemental Figure 6

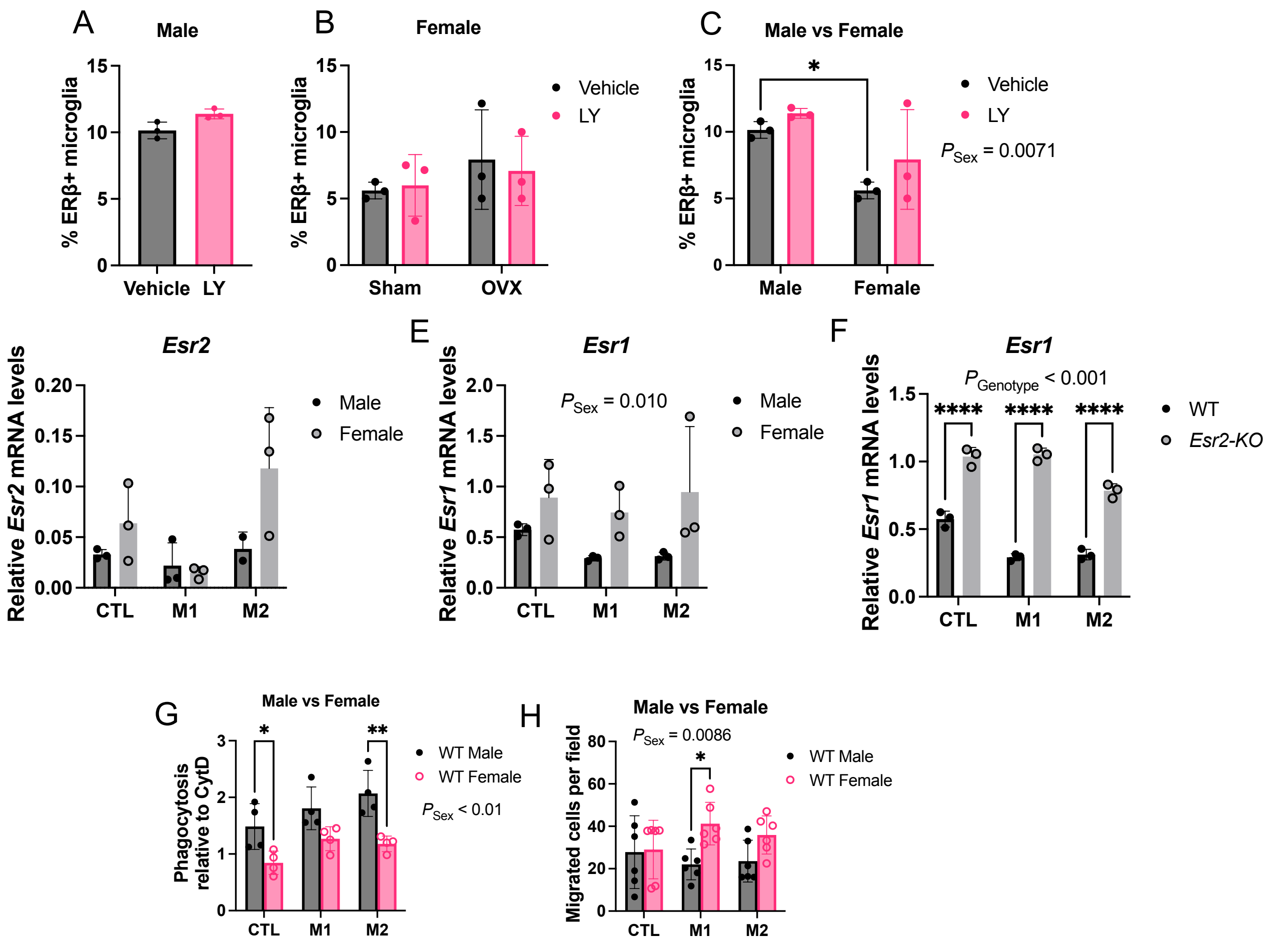


**SUPPLEMENTAL FIGURE 6. ER expression and effect of ERβ knockout (*Esr2*-KO) in male and female mouse microglia.** Percent ERβ positive microglia in male (A) and female (B) $App^{NL\text{-}G\text{-}F}$ brain (cortex and hippocampus) upon OVX and/or LY treatment (n = 3). (C) Comparison of percent ERβ positive microglia between male and female brains. (D) Expression of ERβ (*Esr2*) or (E) ERα (*Esr1*) mRNA relative to *Rplp0* expression in control (CTL), M1, or M2 activated primary male and female microglia (n = 3). (F) Expression of *Esr1* mRNA relative to *Rplp0* expression in CTL, M1, or M2, WT and *Esr2*-KO primary female microglia. (G) Comparison between male and female WT microglia phagocytosis capacity (relative to Cytochaloasin D, CytD) and (H) migration upon CTL, M1 or M2 activation. Statistical significance was determined using unpaired t-test was used (in A) and 2-way ANOVA followed by uncorrected Fisher's LSD test (in B, C) or Šidák's multiple comparisons test (in D-H). Overall significant effects of sex and genotype are indicated. * *P* < 0.05, ** *P* < 0.01, *** *P* < 0.001, **** *P* < 0.0001
